# Dynamics of Spaetzle morphogen shuttling in the *Drosophila* embryo shapes pattern

**DOI:** 10.1101/505925

**Authors:** Neta Rahimi, Inna Averbukh, Shari Carmon, Eyal D. Schejter, Naama Barkai, Ben-Zion Shilo

## Abstract

Establishment of morphogen gradients in the early *Drosophila* embryo is challenged by a diffusible extracellular milieu, and rapid nuclear divisions that occur at the same time. To understand how a sharp gradient is formed within this dynamic environment, we followed the generation of graded nuclear Dorsal (Dl) protein, the hallmark of pattern formation along the dorso-ventral axis, in live embryos. We show that a sharp gradient is formed through extracellular, diffusion-based morphogen shuttling that progresses through several nuclear divisions. Perturbed shuttling in *wntD* mutant embryos results in a flat activation peak and aberrant gastrulation. Re-entry of Dl into the nuclei at each cycle refines the signaling output, by guiding graded accumulation of the *T48* transcript that drives patterned gastrulation. We conclude that diffusion-based ligand shuttling, coupled with dynamic readout, establishes a refined pattern within the diffusible environment of early embryos.

## Introduction

The crude onset and subsequent refinement of spatial information shapes the future body pattern of embryos. Morphogens, key instructive elements in this context, are secreted signaling molecules that induce cells to adapt different fates depending on their concentration. Establishing a morphogen gradient over a field of naïve cells patterns the cell layer into distinct domains of gene expression (Green and Sharpe, 2015; Wolpert, 1971). Different strategies to guide morphogen distribution have been identified. A common option is to produce the morphogen in a restricted group of cells, giving rise to its graded distribution in the surrounding cells (Lecuit et al., 1996; Nellen et al., 1996). Notably, in this scenario, the morphogen-producing cells are distinct from the responding cells.

An alternative strategy of morphogen distribution is applicable to situations where the morphogen is broadly expressed, and the gradient is generated *within* the field of expressing cells, which also respond to the morphogen. This scenario is applicable to early embryos, where broad transcriptional domains have been established, but have not yet given rise to the determination of sufficiently restricted groups of cells, which could provide a local morphogen source. In such situations, restricting morphogen signaling to a narrow domain becomes a challenge, as diffusion tends to spread, rather than restrict ligand distribution.

Studies in several systems identified the *Shuttling* mechanism as a robust solution to this challenge (Shilo et al., 2013). Here, a morphogen gradient is established not merely by its diffusion away from the production source, but through an effective translocation of the morphogen into the center of the field. This translocation, which is purely diffusion driven, is mediated by a proximally-produced inhibitor. The resulting gradient is sharp and robust, displaying limited sensitivity to gene dosages or reaction rate constants. Shuttling provides robustness by concentrating the morphogen into restricted domain, which allows storing excess levels in regions of maximal signaling without modifying the resulting cell fates. Such a shuttling mechanism establishes the bone morphogenetic protein (BMP) morphogen gradient in the early embryos of *Drosophia* and other insects (Eldar et al., 2002; Lapraz et al., 2009; Shimmi et al., 2005; van der Zee et al., 2006; Wotton et al., 2017). It is also used for forming the BMP gradient in the *Xenopus* embryo, where it acquired additional features that allow scaling of the gradient with embryo size (Ben-Zvi et al., 2014; Ben-Zvi et al., 2008).

Compelling evidence for shuttling was provided by comparing mutant phenotypes with the predictions made by computational models (Ben-Zvi et al., 2008; Eldar et al., 2002; Haskel-Ittah et al., 2012). It was also demonstrated that ligand produced ectopically in one part of the embryo can be translocated to and endocytosed in the normal activation domain (Reversade and De Robertis, 2005; Wang and Ferguson, 2005). Experimentally, these data were obtained through the analysis of fixed embryos. Yet, the essence of the shuttling mechanism resides in its *dynamics.* What is the time-frame during which the gradient is established? How fast is gradient formation relative to its readout? Is the gradient stably formed, or is it subject to subsequent cycles of refinements? Insight into these questions requires monitoring the dynamic distribution of the morphogen within single embryos.

Furthermore, the shuttling mechanism makes a number of counter-intuitive predictions regarding the dynamics of pattern formation. In particular, it predicts that signaling at the edge of the source will initially increase, as ligand begins to accumulate, but will subsequently be reduced, since ligand is continuously being shuttled to the center of the field. This nonmonotonic behavior is a defining property of the shuttling mechanism that concentrates ligand, but is absent from other diffusion-based mechanisms establishing a graded pattern. In a certain parameter range, shuttling also predicts transient formation of a double-peak pattern within the gradient, again a prediction that is absent from naïve gradient-forming mechanisms. Uncovering such features is again possible only by monitoring the dynamics of gradient formation in live embryos.

The ability to observe the dynamics of morphogen gradient formation is challenging. The ligands typically function at low levels. Furthermore, the morphogen may be present not only in its active form, but also in a non-processed, inactive form, or bound to an inhibitor. Finally, the morphogen is present in both extra- and intra-cellular locations. Most studies therefore follow the patterning processes by quantifying the intracellular outcome of morphogen signaling as a proxy for active morphogen distribution, using antibodies against the activated (e.g, phosphorylated) states of signaling pathways triggered by morphogens (Dorfman and Shilo, 2001; Gabay et al., 1997; Tanimoto et al., 2000). Such approaches, however, cannot be used to follow live embryos as they rely on immunostaining of fixed samples. An alternative is to follow the transcriptional outcomes of morphogen signaling, but this analysis is already quite removed from the original morphogen gradient itself, and is compounded by additional regulatory inputs controlling the expression of the target genes.

Pattering the dorso-ventral (D-V) axis of the *Drosophila* embryo provides a powerful system to analyze the dynamics of morphogen gradient formation. The early *Drosophila* embryo is a syncytium, that is, a collection of nuclei that occupy a common cytoplasm enclosed by the embryonic plasma membrane. Within this syncytium, the nuclei undergo 13 rapid and synchronous divisions without any change in embryo size or shape. During the final four division cycles, the nuclei form a monolayer just beneath the plasma membrane. Spaetzle (Spz) is secreted from the syncytium into the surrounding peri-vitelline fluid, positioned between the plasma and vitelline membranes, and a gradient of active Spz forms within it, along the radial D-V axis. The processed Spz morphogen binds the transmembrane Toll receptor (DeLotto and DeLotto, 1998; Morisato and Anderson, 1994; Schneider et al., 1994; Weber et al., 2003), and triggers Dorsal (Dl) translocation into the syncytial nuclei (Figure 1A). The processed form of the Spz ligand therefore functions as the morphogen at this stage.

**Figure 1.**
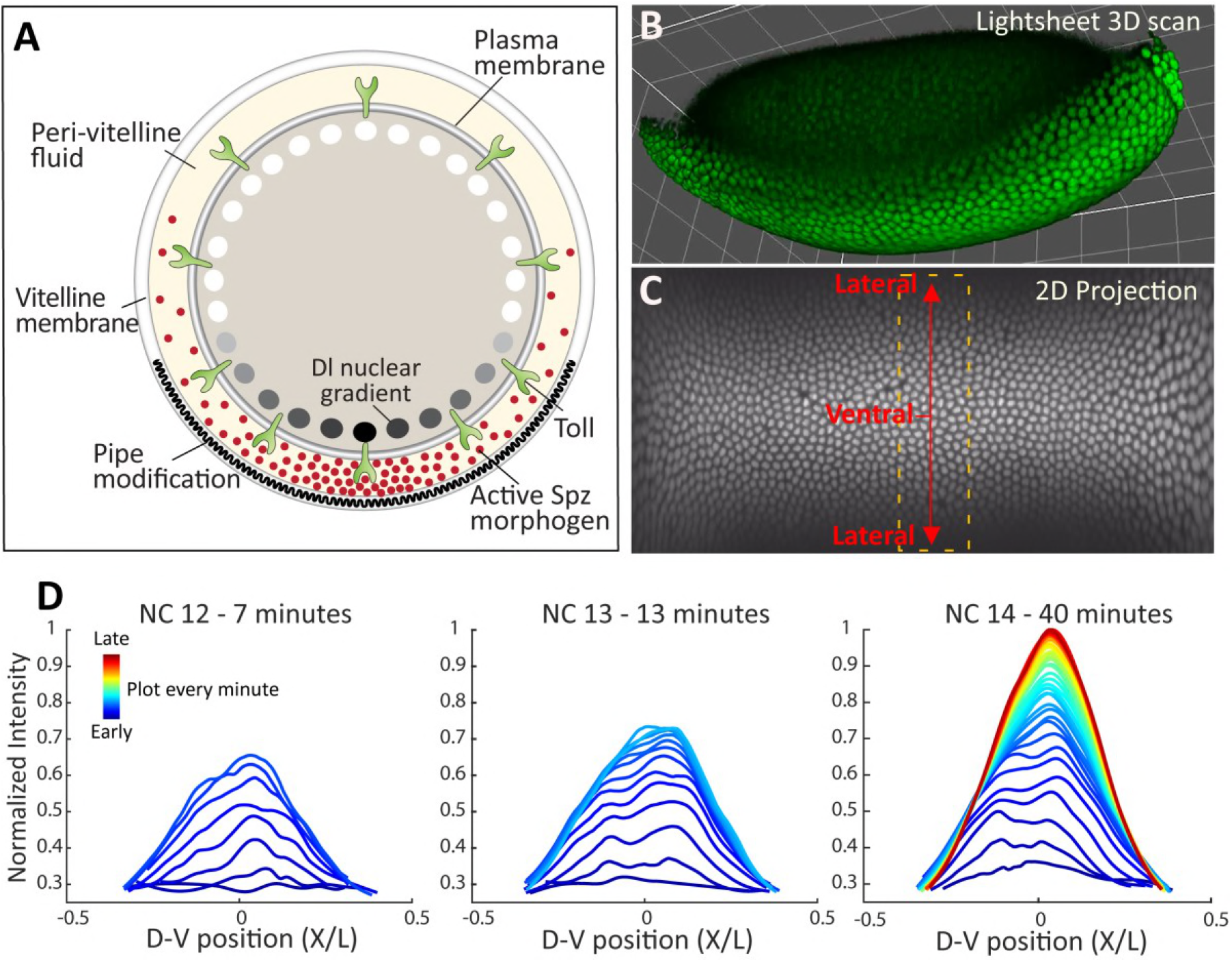
Dynamics of Dl-nuclear localization. (A) Schematic cross-section of an early embryo: Spz is distributed in a sharp gradient along the D-V axis within the peri-vitaline space. Spz binding to the Toll receptor triggers a signaling cascade, culminating in a gradient of nuclear Dl with a sharp ventral peak. Generation of the Spz sharp gradient was proposed to employ a self-organized shuttling mechanism (Haskel-Ittah et al., 2012). (B) A frame from a Light Sheet time-lapse movie following the dynamics of endogenously expressed Dl-GFP in the entire embryo. The gradient can be visualized already at NC 12, it is lost during nuclear divisions and is re-generated at the onset of each nuclear cycle. (C) Each frame of the 3D Light Sheet scan was projected to a 2D map using the ImSAnE tool (Heemskerk and Streichan, 2015). Here, the 2D projection of the frame from B is shown. The quantification of Dl-GFP intensity in the nuclei along the D-V axis at 1 min intervals was carried out on nuclei inside the dashed frame. This region, which is close to the middle of the A-P axis, is not distorted by the 2D projection (See Methods). (D) Dl-GFP intensity, plotted as function of relative location along the D-V axis (relative location axis x/L is defined as location divided by embryo circumference (See Methods)). A curve is shown for each time point for NC 12, 13 and 14 for the same *wt* embryo. Each Dl-GFP intensity curve was smoothened and normalized by the maximal value attained during NC14 (See Methods). Time points are one minute apart, going from earliest time points in blue to the latest in red. Duration of each NC in minutes is indicated above the plot.

We previously showed that graded active Spz distribution is established by a shuttling mechanism. In this case, shuttling is implemented in a self-organized manner through a complex interplay between the active ligand and its pro-domain, which can accommodate diverse structures (Haskel-Ittah et al., 2012; Shilo et al., 2013). The resulting gradient of active Spz is sharp, and provides robustness to a variety of perturbations in the level of pathway components.

Entry of Dl into the nuclei can be followed in single live embryos carrying a Dl-GFP fused protein (DeLotto et al., 2007). In this work, we use Light Sheet fluorescence microscopy for live imaging of Dl-GFP nuclear localization during the final nuclear division cycles of the syncytial *Drosophila* embryo. The resulting dynamics shows the two signatures of ligand shuttling: a transient increase in signaling in the lateral regions, which is then reduced so as to preferentially increase signaling at the ventral midline, and the resolution of two lateral peaks to a single central peak. We find that ligand shuttling is an ongoing process, which repeats itself following each nuclear division. During the beginning of nuclear cycle (NC) 14, the resulting dynamics of nuclear re-entry of Dl allows to further refine the resulting spatial pattern, by triggering different temporal onsets of zygotic target gene expression in closely positioned nuclei, thereby leading to a functionally significant graded accumulation of target gene transcripts. In *wntD* mutant embryos, the Dl peak becomes flattened, and leads to an abnormal increase in the number of cells simultaneously undergoing the initial step of gastrulation, underscoring the significance of timely and properly shaped gradient formation. Thus, diffusion-based ligand shuttling, coupled with a dynamic readout, establishes a refined pattern within the environment of early embryos.

## Results

### Temporal evolution of the Spz gradient during nuclear cycles 12-14

Using Light Sheet fluorescence microscopy, we followed individual embryos carrying a Dl-GFP fusion protein expressed under the endogenous *Dl* promoter (Figure 1B, SI: movie 1). Consistent with previous reports (DeLotto et al., 2007; Kanodia et al., 2009), we observed a D-V gradient of nuclear Dl-GFP already at NC 12. This gradient was further refined and elaborated during the next two cycles. To enable quantitative analysis of Dl-GPF nuclear dynamics, we used an area preserving transformation to project the 3D images onto a 2D sheet. We restricted our analysis to a region surrounding the A-P midline, where distortion due to 2D projection is negligible (Figure 1C, SI: movie 2) (Heemskerk and Streichan, 2015). Next, we automatically segmented the nuclei and averaged the nuclear Dl-GFP signal in nuclei occupying a similar D-V axis position.

Our measurements defined the quantitative, spatio-temporal dynamics of Dl-GFP at a 1-2 minute time resolution (Figure 1C-D, SI: movie 3, Methods). This dynamics results from the extracellular active Spz gradient. However, inferring the profile of this extracellular gradient from Dl-GFP dynamics is confounded by the fact that Dl-nuclear accumulation is established anew at every nuclear cycle, since Dl exits the nucleus at mitosis upon nuclear envelope breakdown. We therefore needed a framework to suitably infer properties of the extracellular active Spz gradient, and critically distinguish between models of gradient formation.

Toll signaling, at each given position along the D-V gradient, triggers the level of nuclear Dl and the rate by which this level increases. Thus, at the beginning of each division cycle, following re-establishment of the nuclear envelope, nuclear Dl levels increase at a rate that is proportional to the level of nearby Toll signaling. Conversely, at longer times, nuclear Dl levels approach a steady state, and are proportional to the extracellular Toll signaling. We therefore plotted the dynamics of both parameters, nuclear Dl levels and its temporal change during the onset of NC 14. Notably, we observe that this qualitative dynamics differs, depending on the spatial position of nuclei along the D-V axis. In the ventral-most regions they increased monotonically. In contrast, in lateral domains nuclear Dl displayed an overshoot, initially increasing but then starting to decrease (Figure 2A-D, Figure S1). Clearly, such a decrease in nuclear Dl is only possible if Toll signaling at this position decreases as well. Therefore, the data indicates that the external Spz gradient continues to evolve through the early part of NC 14, showing a distinct position-dependent, non-monotonic temporal signature.

**Figure 2.**
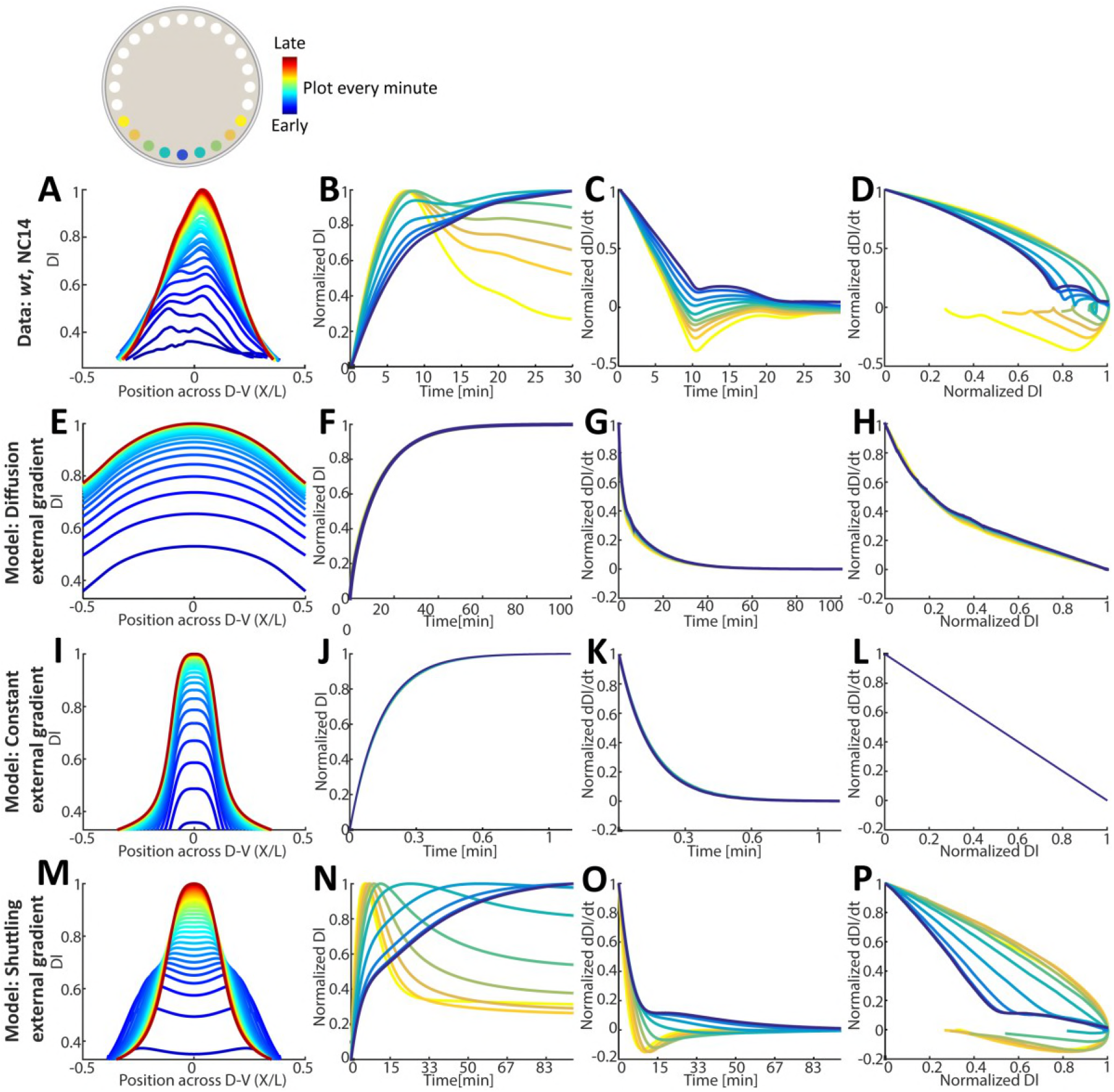
Measured nuclear Dl dynamics during NC14 indicates ongoing shuttling. (A-D) Measured nuclear Dl temporal dynamics for a *wt* embryo, time points are color coded and are one minute apart. (E-P) Simulations of nuclear Dl temporal dynamics (See SI). (E-H) Diffusion model: nuclear Dl localization induced by Spz binding the Toll receptor, with Spz diffusing out of a wide source while being degraded. (I-L) Constant external gradient model: nuclear Dl localization induced by Spz binding the Toll receptor, with a Spz gradient that is constant in time. (M-P) Full model: nuclear Dl localization induced by Spz binding the Toll receptor, with shuttling of Spz. (A,E,I,M) Nuclear Dl levels in [a.u], plotted as function of relative location along the D-V axis. A curve is shown for each time point, time points are color coded. (B,F,J,N) Nuclear Dl levels at specific locations along the D-V axis, as function of time. Nuclear localizations along the D-V axis are color coded. Each location’s curve was normalized by its own maximal value. (C,G,K,O) Nuclear Dl temporal derivative at specific locations along the D-V axis, as function of time. Each location’s curve was normalized by its own maximal value. (D,H,L,P) Nuclear Dl temporal derivative at specific locations along the D-V axis, as function of nuclear Dl. Each location’s curve was normalized by its own maximal value. Color codes for the location of nuclei along the D-V axis and for the temporal dynamics are shown.

To more rigorously infer dynamic properties of the external gradient from the measured pattern of nuclear Dl, we used computer simulations, modeling Dl-nuclear entry while assuming different temporal patterns of Toll signaling (Figure 2E-L). Specifically, we searched for a qualitative signature that distinguishes between three scenarios: (1) constant Toll signaling; (2) Toll signaling that is changing (increasing) monotonically in time, as expected in naïve gradient-forming models; and (3) a non-monotonic increase in Toll signaling, the signature found in lateral regions of gradients formed by the shuttling mechanism. Our simulations have shown that these scenarios are best distinguished by comparing the temporal changes in nuclear Dl (d(Dl)/dt) with the levels of nuclear Dl. In the first two cases – constant or monotonically increasing Toll activity – the relation between these two parameters is invariably linear or concave (Figure 2E-L). In contrast, in the presence of non-monotonic shuttling-based dynamics, a convex relation is obtained, with a pronounced negative temporal derivative at the lateral regions, where nuclear Dl levels are low (Figure 2M-P).

The measured data is not consistent with the dynamic defined by constant, or monotonically increasing Toll signaling. Rather, it shows a clear signature of non-monotonic, shuttling-like dynamics. Extending our simulations to include the full shuttling model that establishes the active Spz gradient combined with Dl nuclear transport (See SI), confirmed that this model is fully capable of simulating the experimentally observed dynamics, including the nonmonotonic, overshoot dynamics at the lateral regions.

An additional notable property of Dl-nuclear entry dynamics was the initial formation, at every nuclear cycle, of two ventro-lateral signaling peaks, that eventually converge to a single ventral peak (Figures 1D, 2A, Figure S1). Thus, by 10-15 minutes into NC 14, when the major target genes for Dl are induced, the initial two-peak gradient has refined to a single sharp peak. The initial two-peak pattern provides another unique signature of shuttling-like dynamics. It is expected under certain shuttling parameters, when the mean distance traveled by the shuttling complex before it is cleaved, is much smaller than the distance to the ventral-most site. In this case, ligand will initially accumulate at lateral regions, followed by gradual ventral translocation (See SI). The reappearance of the double peak at every nuclear cycle likely reflects a process of extracellular ligand mixing in the peri-vitelline fluid, possibly caused by reorganization of the cortical actin-based cytoskeleton and deformation of the plasma membrane associated with the nuclear divisions (di Pietro and Bellaiche, 2018; Zhang et al., 2018).

In conclusion, the dynamic behavior of Dl-GFP supports a continuous process of extracellular Spz shuttling, displaying two of its defining signatures: non-monotonic dynamics of nuclear Dl entry in lateral regions, and the transient formation of two-peak gradient.

### Altered shuttling dynamics in *wntD* mutants affects gastrulationg

Dl-nuclear localization dynamics can be used for refined analysis of informative mutant phenotypes. We applied this approach to study WntD, an inhibitor of Toll signaling which provides a negative feedback that buffers the D-V patterning gradient against fluctuations (Rahimi et al., 2016). *wntD,* a target of the Toll pathway, is transcribed locally at the posterior terminus of the embryo, and the secreted protein diffuses within the peri-vitelline fluid to attenuate Toll signaling (Figure 3A)(Helman et al., 2012).

**Figure 3.**
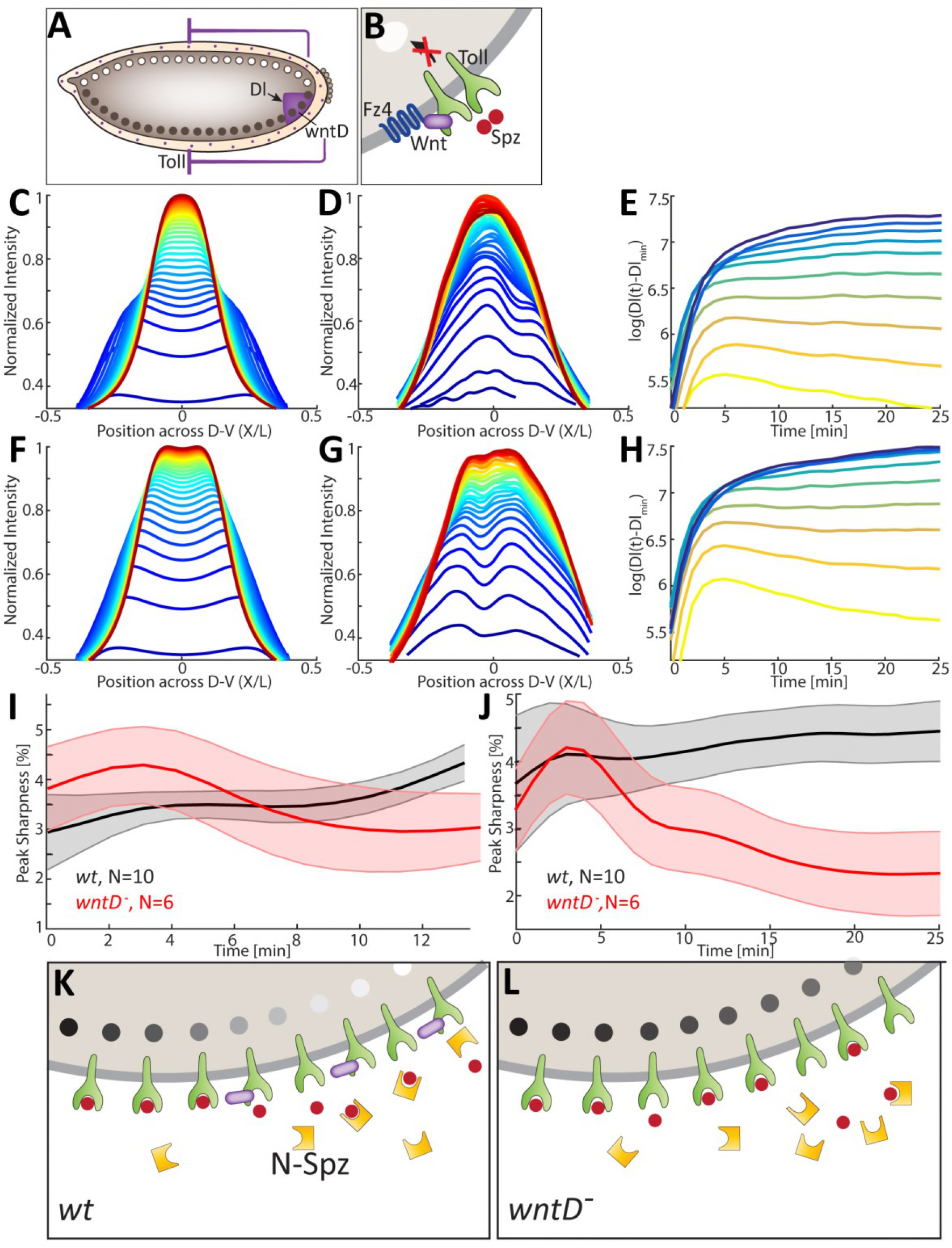
Shuttling dynamics are impaired in the absence of WntD. (A) A scheme showing the integral feed-back loop between Dl and WntD. *wntD* expression is restricted to the posterior side of the embryo (purple) and is induced by Dl. (B) A scheme showing WntD binding to Fz4, and restricting the binding of Spz to the Toll receptor. (C,F) Simulations of Dl temporal dynamics in the full model for a *wt* (C) and *wntD* mutant (F) embryo (See SI). Time points are color coded. For the *wntD* mutant, the double peak is more prominent and does not fully converge resulting in a flat peak. (D,G) Measured Dl temporal dynamics for a *wt* (D) and *wntD* mutant (G) embryo (See Methods). Time points are color coded, and are one minute apart. As predicted by the model, the *wntD* mutant exhibits less efficient shuttling leading to a more prominent double peak, which does not fully converge, resulting in a flat peak. (E,H) Log of Dl-GFP intensity temporal dynamics at selected locations along the D-V axis for a *wt* (E) and *wntD* mutant (H) embryo from panels D,G. Locations are color coded. Measured values were background subtracted and smoothed in time (See Methods). The flat peak of the *wntD* mutant embryo results in very similar Dl values over time for the four ventral most curves, while in the *wt* embryo each curve attains a different final value and has different dynamics. (I-J) Peak sharpness over time (calculated as std/mean in % of values close to the peak, See Methods) in a population of *wt* embryos (black) and *wntD* mutant embryos (red) during NC 13 (I) and NC14 (J). Bold lines indicate population mean and surrounding color indicates standard error. Number of embryos in each population is indicted in plot. Sharpness in the *wt* population increases over time to a value significantly higher than the *wntD* mutant population. The *wntD* mutant population exhibits an initial increase in peak sharpness due to the prominent double peak, followed by a decrease due to peak flattening. (K) Scheme showing the global attenuation effect of WntD on Toll availability, leading to redirection of active ligand binding towards more ventral positions. (L) In *wntD* mutants additional Toll receptors become available, leading to increased ligand binding in lateral regions, generating a flatter peak of signaling.

Genetic epistasis experiments have demonstrated that WntD binds the Frizzled-4 (Fz4) receptor, and associates with the extracellular domain of Toll (Rahimi et al., 2016)(Figure 3B). However, the consequences of this inhibition on gradient formation remained unclear. One option is that WntD uniformly decreases signaling in all regions. Alternatively, with the shuttling mechanism in mind, our simulations suggested that the binding of WntD to the extracellular domain of Toll promotes ligand shuttling. This is because the binding of WntD to the Toll receptor decreases the number of available receptors. This, in turn, compromises the ability of free Spz to bind Toll, thereby increasing the probability that it will bind the free shuttling molecule. Binding of Spz to Toll will thus be directed to more ventral regions, where the levels of the shuttling molecules are sufficiently low. Uniform distribution of moderate WntD levels would therefore lead to redistribution of the ligand to more ventral regions, impacting not only on the strength of Toll signaling, but also on its sharpness (see Figure 3C,F for simulation results).

Using live imaging in a *wntD* mutant background, we directly tested the possibility that WntD promotes shuttling. Indeed, in *wntD* mutant embryos shuttling was less efficient. First, the initial signature of a double peak was more prominent and persistent. Second, in contrast to *wt* embryos where gradual sharpening of the gradient takes place, in a wntD-mutant background the final gradient was relatively wide and displayed a flattened single peak (Figures 3C-J, S2) – consistent with the predicted role of WntD in redistribution of the Spz ligand (Figure 3K-L).

To follow the morphological consequences of a wider peak distribution of nuclear Dl distribution, we monitored *wntD* mutant embryos for an extended period of NC 14, observing the processes of gastrulation and ventral furrow formation. We defined the edges of the furrowing domain by marking the two lateral-most nuclei that alter their orientation upon gastrulation. Working backwards to an earlier phase of NC 14, when the nuclei are still in a monolayer, we can accurately count the number of nuclei between these edges. In contrast to gastrulating *wt* embryos where the initial invagination is observed in ~9 cells, in *wntD* mutants a broader front of up to 15 cells invaginated at the same time (Figure 4). Thus, the shape of the Dl-activation gradient is essential for normal patterning and gastrulation. When the final gradient peak is not sharp, a larger cohort of ventral cells takes part in furrow formation.

**Figure 4.**
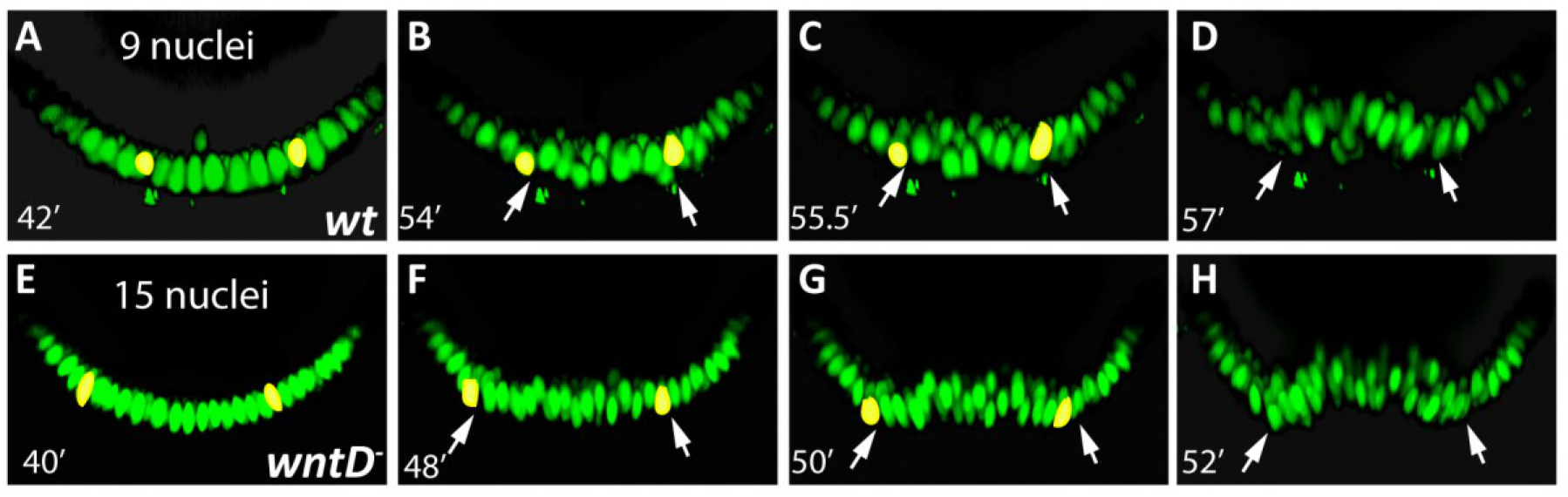
An enlarged ventral furrow is formed in the absence of WntD. Snapshots of cross-sections of a *wt* (A-D) vs. *wntD* mutant embryo (E-F) expressing Dl-GFP, showing the ventral nuclei during NC 14. Time in NC 14 (in minutes) is indicated. When the first sign of invagination appeared, the most lateral nuclei still displaying positional change were marked (yellow). Working backwards in the movies allowed an accurate count of the nuclei between them prior to invagination.

### Timing of *wntD* transcription

*wntD* mutants display perturbed Dl dynamics already at NC 13 (Figures 3H, S3) suggesting that WntD normally exerts its modulating effects at this early stage. This implies that zygotic expression of *wntD,* its translation, secretion to the peri-vitelline fluid and diffusion of the protein, have commenced by then. To examine that this is possible, we applied our Light Sheet-based visualization setup as a more sensitive assay for defining the onset and temporal dynamics of *wntD* expression.

*wntD* expression is activated by nuclear Dl, and is restricted to the posterior region, where Torso signaling relieves Capicua (Cic) repression (Helman et al., 2012). We generated a wntD::MS2 reporter, utilizing the genomic upstream regulatory sequence of *wntD* (Figure 5A). The early expression profile of the reporter at the posterior part of the embryo mimics the known pattern of *wntD* (Rahimi et al., 2016). Importantly, all eight embryos examined expressed *wntD,* implying that overshooting of Toll signaling, which triggers expression of the WntD “buffer”, is a common consequence of D-V gradient signaling. When monitoring the dynamics of *wntD* expression, the number of active nuclei transcribing *wntD* decreased continuously between NCs 12 and 14 (Figure 5B-F). These results suggest that the WntD protein already exerts its attenuating effect on Toll signaling at an early time.

**Figure 5.**
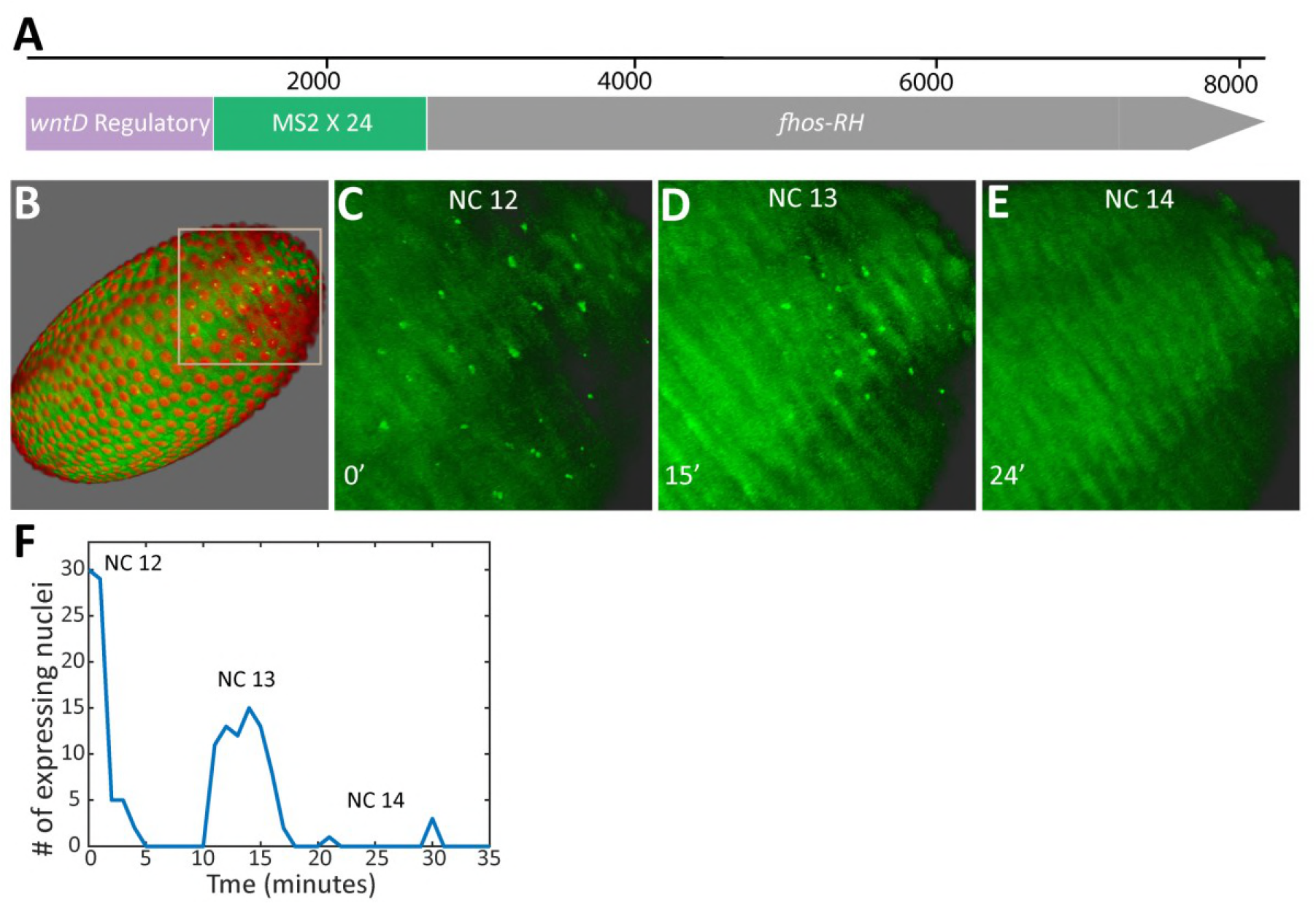
Dynamics of *wntD* transcription. (A) Schematic representation of the wntD::MS2 reporter. The 1.2 kb regulatory region upstream of the *wntD* coding sequence drives the expression of Fhos-RH with 24 MS2 repeats located at its 5’ end. (B) Light Sheet live imaging of the transcriptional activity of the *wntD* promoter enables a quantitative analysis of its dynamics. Embryos express His:RFP which marks nuclei in red and the transcriptional activity is indicated by the green signal. (B-F) A constant reduction in the number of nuclei expressing *wntD* is observed from NC12 onwards, such that by NC 14 no transcription is observed. Since the *wntD* promoter responds to the level of Dl, we assume that this reduction reflects the attenuating activity of WntD.

### Dl nuclear re-entry promotes graded expression of the zygotic *T48* gene by different temporal onsets of transcription

The gradient of Dl-nuclear localization defines three major domains of zygotic gene expression along the D-V axis. Within each of these domains several target genes are uniformly expressed. The mesoderm is defined by highest levels of nuclear Dl and uniform expression of the zygotic target genes *twist (twi)* and *snail (sna)* (Rusch and Levine, 1996). However, a graded zygotic response within the mesoderm is also required: A gradient of apical myosin II recruitment, peaking at the ventral midline, is essential for the ordered apical cell constriction driving ventral furrow formation (Heer et al., 2017). A zygotic target gene that may lead to graded myosin II distribution is *T48,* since the T48 protein recruits Rho GEF2 to the apical membrane, triggering the accumulation and contractile activity of an apical actomyosin network (Kolsch et al., 2007). We asked whether the dynamics of Dl-nuclear entry may play a role in generating a graded transcriptional response within this region, by following the zygotic target gene *T48.*

Previous analysis of an MS2 reporter for *T48* transcription demonstrated that the ventral-most nuclei initiate transcription earlier than the lateral ones (Lim et al., 2017). Using the same reporter, we find that signal intensity in transcribing nuclei is similar regardless of their position along the D-V axis, suggesting that once *T48* transcription is initiated, it progresses at a constant rate in all nuclei (Figure 6C). If zygotic expression of Dl-target genes depends not only on the final, steady-state level of nuclear Dl, but also on the dynamic profile of its accumulation, the signaling output could be further sharpened. A dynamic phase of Dl nuclear entry takes place during the initial 20 minutes of NC 14 (Figure 1D), the nuclear cycle associated with a major onset of zygotic gene expression. A consequence of these dynamics is that ventral-most nuclei will reach the threshold for expression of a given zygotic gene earlier than more lateral ones. These ventral nuclei will begin to express the gene earlier, therefore expressing it for a longer period than more lateral nuclei, and could thus accumulate more transcripts.

**Figure 6.**
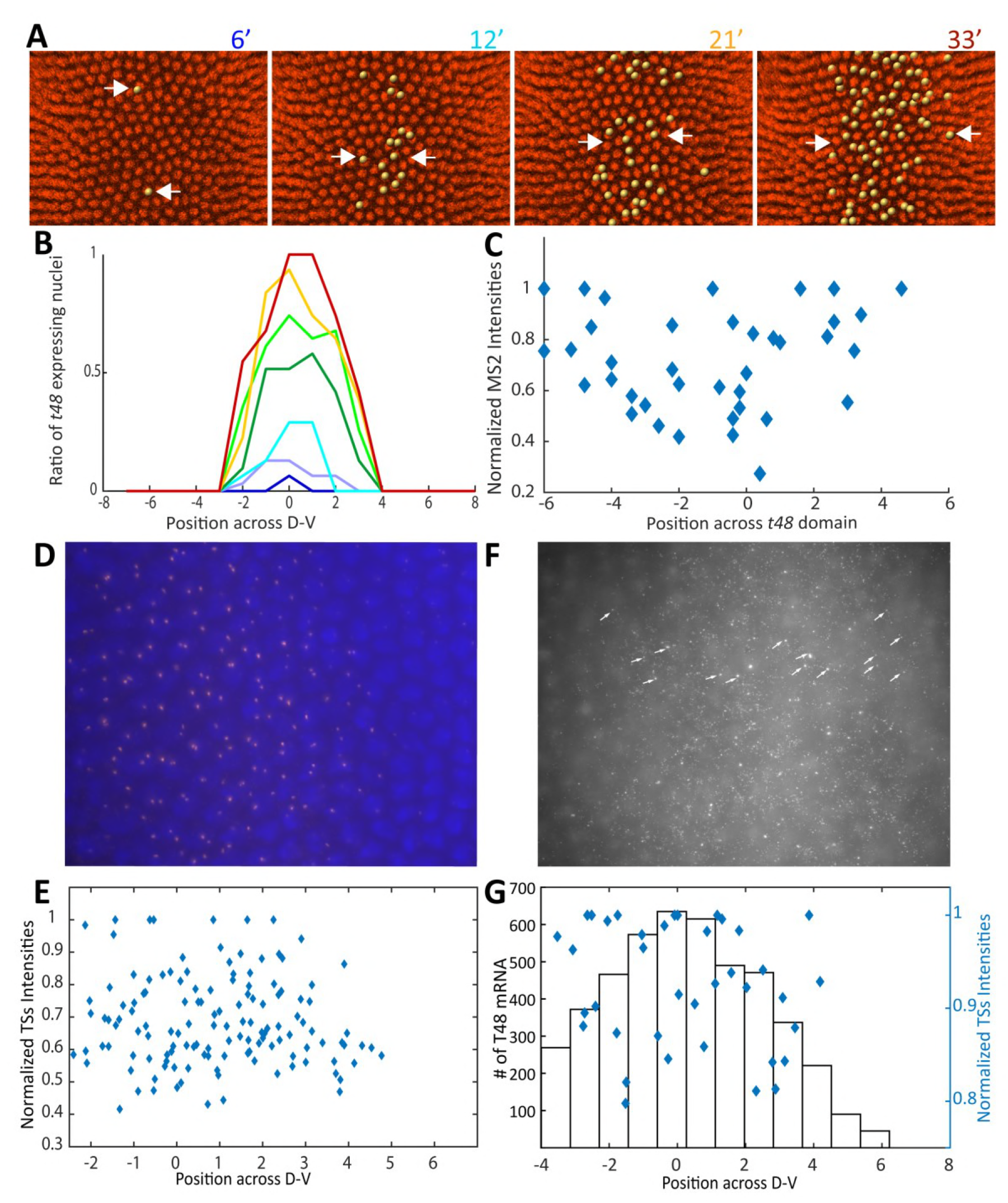
Graded accumulation of T48 transcripts is driven by the dynamics of Dl-nuclear entry. (A) Time lapse images from the beginning of NC 14, showing the dynamics of the onset of *T48* transcription in a *wt* embryo, followed by T48::MS2. Nuclei displaying transcription were highlighted in yellow. Arrows mark the edges of the domain at each time point. (B) Quantification of the data in (A). Times are color-coded according to (A), Y axis represents the normalized number of transcription start sites (TSs) along the A-P axis, X axis represents the position along the D-V axis. (C) The intensity of T48::MS2 signal for nuclei across the D-V axis at 33 min. Once *T48* transcription is initiated, it progresses at comparable rates in all nuclei. (D,E) Single molecule FISH for *T48* showing the intensity of transcription and the number of active nuclei across the region expressing *T48.* Nuclei are marked by DAPI (blue) and *T48* probe in red. Once transcription is initiated it proceeds at a similar rate by all nuclei, and the number of nuclei expressing *T48* in ventral and ventro-lateral regions is comparable. (F) The ventral *T48* expression domain, arrows mark TSs. (G) Quantification of smFISH data: Number of *T48* mRNA molecules in black bars, and TSs intensity corresponding to transcription rate in blue diamonds.

To examine the consequences of the graded onset of *T48* transcription on mRNA accumulation, we carried out quantitative single-molecule FISH using *T48* probes. The signal obtained is comprised of two components: Prominent puncta in the nuclei representing active transcription of the gene, and sparse weaker spots in the cytoplasm marking accumulated individual mRNA molecules. We analyzed embryos that demonstrated ongoing transcription in several nuclear rows along the D-V axis. The number of transcribing nuclei in ventral and lateral positions within the expression domain was similar, and the transcription intensity in the different rows appears comparable. This again indicates that once *T48* transcription is initiated, it progresses at a constant rate in all nuclei, regardless of position (Figure 6D-E). We next quantitated the levels of cytoplasmic *T48* mRNA, using the TransQuant (Bahar Halpern and Itzkovitz, 2016). A clear D-V gradient of cytoplasmic mRNA accumulation is observed across several cell rows, peaking at the ventral midline (Figure 6F-G). This result indicates that the *T48* mRNA is sufficiently stable during the temporal window of early NC 14, such that the time of onset of its transcription along the D-V axis, governed by the dynamics of Dl nuclear entry, correlates with the level of mRNA that accumulates in the adjacent cytoplasm. Dl nuclear entry dynamics thus appear to be a critical factor regulating graded *T48* activity along the D-V axis.

## Discussion

### Dynamics of the Spz extracellular morphogen gradient

The early *Drosophila* embryo provides extreme challenges for the generation and maintenance of extracellular morphogen gradients. Most notably, the peri-vitelline fluid surrounding the embryo facilitates rapid diffusion of molecules (Stein et al., 1991). In addition, the alteration in the surface of the plasma membrane at every nuclear division provides an active mixing force (di Pietro and Bellaiche, 2018; Zhang et al., 2018). Thus, analysis of the early morphogen gradients operating in this environment, including ventral Spz/Toll activation and the subsequent BMP gradient patterning the dorsal aspect, should consider this highly dynamic environment. In the case of the Toll pathway, the active Spz ligand is generated by proteolytic processing within the extra-embryonic peri-vitelline fluid in a broad ventral region, defined by the activation domain of the Easter (Ea) protease (Cho et al., 2012). The generation of a sharp Spz activation gradient within this broad ventral domain of processing takes place by diffusion-based shuttling. Our previous work demonstrated that the pro-domain of Spz plays an instructive role in delivering the active, cleaved ligand towards the ventral midline (Haskel-Ittah et al., 2012). While a variety of experiments and computational analyses indicated the utilization of a “self-organized shuttling” mechanism in this context, it was imperative to visualize the actual dynamics of the process.

We were able to infer the dynamics of the extracellular Spz gradient by following the kinetics of Dl-GFP nuclear accumulation in individual live embryos during the final syncytial nuclear division cycles and the early phase of NC 14. Nuclear levels of Dl are not a direct readout of the extracellular gradient, since accumulation of Dl in the nuclei is re-initiated at the onset of every nuclear cycle. Nevertheless, it is possible to infer key features of the extracellular Spz gradient from this dynamic behavior. Using this approach we identified clear hallmarks of ligand shuttling, most notably the lateral overshoot and the presence of two lateral peaks which converge to a central ventral peak. This convergence takes place within a timeframe of minutes, and repeats at every nuclear cycle. Since new protein molecules of the extracellular components are continually translated, the ongoing activity of the shuttling process is vital. Therefore, shuttling is important not only for generating the gradient, but also for maintaining it, in the face of rapid diffusion and mixing within the peri-vitelline fluid. Importantly, by ~10-15 minutes into NC 14, when the robust induction of transcription of the cardinal zygotic Dl-target genes *twt* and *sna* ensues, the nuclear gradient of Dl is sharp and a single ventral peak is resolved.

### The role of WntD in shaping and buffering the Spz gradient

Having described the dynamics of Dl-nuclear entry and gradient formation, we were in a position to use our experimental approach in order to examine regulatory processes affecting Toll signaling. The Wnt family ligand WntD provides an essential buffering system to variations in Toll signaling between embryos (Rahimi et al., 2016). *wntD* is an early zygotic gene that is expressed initially at the posterior-ventral region of the embryo, and its expression levels depend on the magnitude of Toll signaling (Helman et al., 2012). Although WntD is produced locally, the rapid secretion and diffusion of the protein in the peri-vitelline space generates a uniform attenuation of Toll signaling throughout the embryo surface. The activity of WntD leads in different embryos to convergence of the variable global Toll activation gradient to a similar pattern, which is dictated by the fixed final signaling level that shuts off *wntD* expression (Rahimi et al., 2016). We term this paradigm “distal pinning”, achieved in this case by an induction-contraction mechanism (Shilo and Barkai, 2017).

Secreted WntD is recruited to the plasma membrane by binding to its receptor Fz4. Epistasis assays have indicated that WntD exerts its inhibitory effect on Toll signaling by associating with the extracellular domain of Toll (Rahimi et al., 2016), thereby reducing the number of Toll receptors that are available for binding Spz. Bearing the cardinal features of shuttling in mind, this mode of inhibition implies that the effect of WntD would be global and non-autonomous, and will actually change the shape of the gradient, making it sharper. The observed dynamics of Dl-GPF in *wntD* mutant embryos indeed confirms this prediction.

The shuttling process is driven by competition between the inhibitory Spz pro-domain and the Toll receptor for binding free, active Spz. Binding to the pro-domain is favored in the lateral part of the embryo, where its concentration is higher, while in more ventral regions binding to Toll takes over. Since WntD impinges on the extracellular properties of the Toll receptor, the active ligand is deposited in more ventral regions, where the concentration of the pro-domain is lower. Thus, WntD does not simply reduce the overall profile of Toll activation, but actually *re-directs* the ligand from the lateral regions to the ventral domain. We have previously shown that accumulation of excess ligand in the peak by shuttling is an effective mechanism to buffer noise. Since activation in this region is already maximal, the excess ligand will not alter the resulting cell fates (Barkai and Shilo, 2009).

The rapid timing of processes in the early embryo and the short duration of interphases between nuclear divisions raises the question of whether it is actually possible to produce sufficient levels of WntD that will drive the morphogen profile to the desired equilibrium. When monitoring *wntD* transcription directly utilizing the MS2 system, we saw that most, if not all embryos express *wntD,* indicating that Toll signaling overshoots in most embryos. Furthermore, within single embryos the number of nuclei expressing *wntD* was reduced between NCs 12 and 13, and completely terminated by NC 14, implying that WntD impinges on the Toll gradient and its own expression by this time. The intronless arrangement of the *wntD* gene and the rapid secretion of the protein, which does not require post-translational modifications (Herr et al., 2012), may facilitate the process.

### Dl nuclear re-entry refines its signaling output

The ventral cohort of zygotic target genes including *twi* and *sna* is induced by the Toll activation gradient, and the threshold for their induction corresponds to ~50% of maximal Dl-nuclear localization (Kanodia et al., 2009; Liberman et al., 2009). Within the ventral domain, nuclei exhibit a similar level of *sna* transcription (Bothma et al., 2015; Lagha et al., 2013). These genes are triggered at NC 14 after the Dl gradient is stabilized and a distinct activation peak generated.

Are there zygotic target genes that respond to the dynamics of Dl nuclear targeting, before it stabilizes? This appears to be the case for *T48*, which encodes a transmembrane protein that facilitates recruitment of RhoGEF2 and ultimately Rho and actomyosin, to mediate apical constriction of invaginating mesodermal cells (Kolsch et al., 2007). Graded distribution of myosin II was shown to be critical for proper invagination of these cells to form the ventral furrow (Heer et al., 2017).

We provide evidence that the graded distribution of *T48* mRNA results from the dynamics of Dl-nuclear re-entry at NC 14. The ventral-most cells reach the threshold of *T48* induction earlier than more lateral cells, and hence will express the gene longer (Lim et al., 2017). Integration of the length of expression along the D-V axis then leads to a gradient of cytoplasmic *T48* mRNA accumulation. This example represents a unique case, where graded morphogen activation instructs the generation of a gradient of target-gene expression. The strict dependence on the *timing* of transcription initiation provides another mechanism to generate differences between adjacent nuclei along the D-V axis.

In conclusion, this work has utilized live imaging of Toll pathway activation, to identify and characterize the hallmarks of ligand shuttling (Figure 7). This process is rapid and takes place continuously throughout the final nuclear division cycles, to generate and maintain a sharp activation gradient in the diffusible environment of the peri-vitelline fluid. WntD impinges on Spz shuttling, and is responsible not only for buffering variability between embryos, but also for generating a sharp activation peak. This peak is utilized to induce a graded expression of a zygotic target gene that is essential for executing processes that drive gastrulation. Thus, diffusion-based ligand shuttling, coupled with a dynamic readout, establishes a refined pattern within the environment of early embryos.

**Figure 7.**
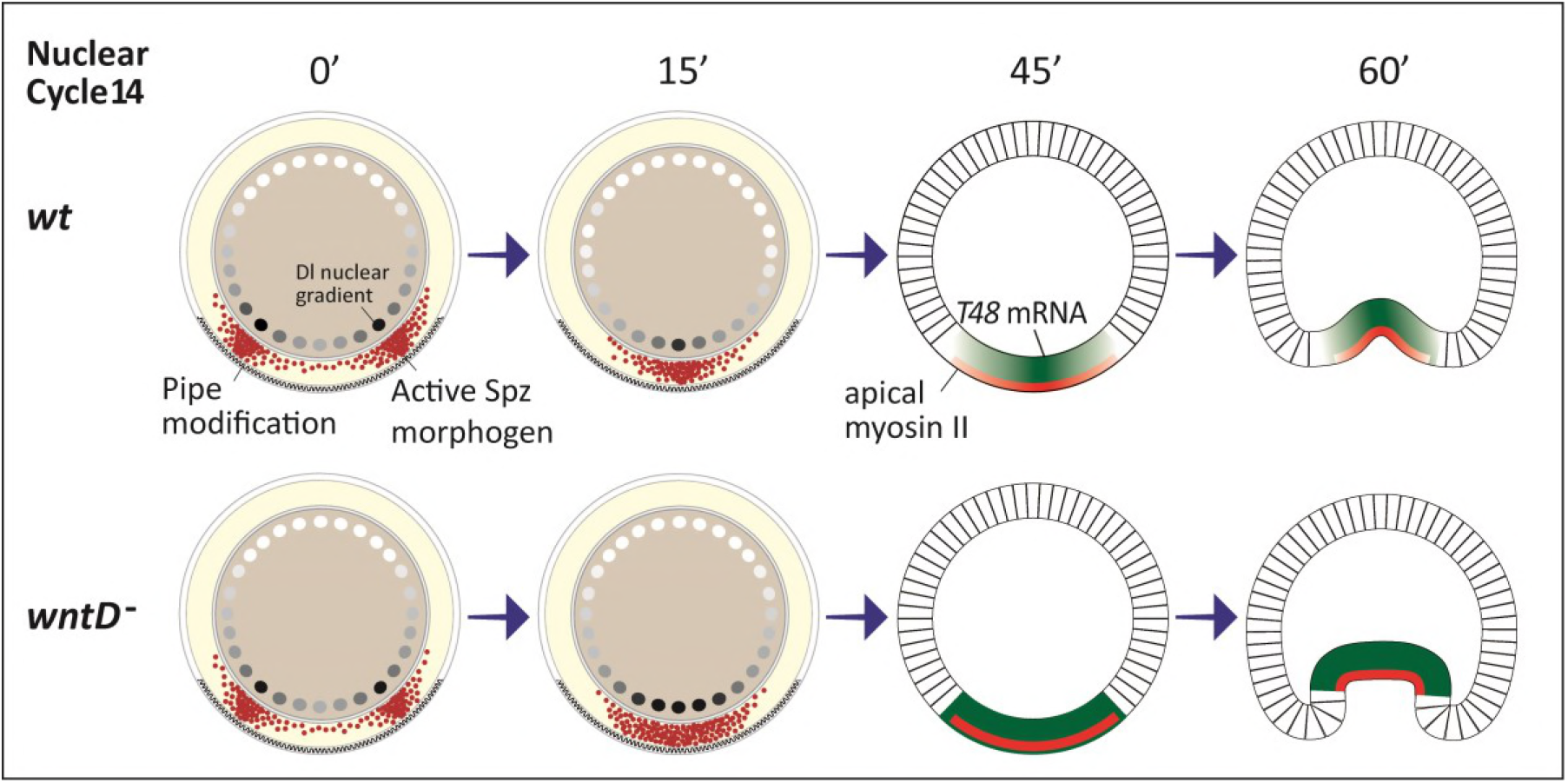
Dynamics of Spz shuttling establishes pattern. The shuttling mechanism operates within a wide domain D-V predefined by Pipe expression during oogenesis. Initially, a doublepeak distribution of the Spz morphogen in the peri-vitelline space is observed. In *wt* embryos shuttling operates efficiently to concentrate the Spz morphogen to a single sharp peak. The dynamics of the Dl gradient are utilized by the embryo to induce graded accumulation of *T48* transcripts, which facilitate recruitment of RhoGEF2 and ultimately Rho1 and actomyosin, to drive apical constriction of invaginating mesodermal cells. Loss of WntD renders shuttling less efficient, leading to a flattened peak of morphogen distribution. This may result in impaired distribution of *T48* transcripts and therefore in a broader gastrulation furrow.

## Acknowledgements

We thank B. Lim and M. Levine for the T48::MS2 flies, and S. Itskovitz for advice on singlemolecule FISH. We are grateful to Y. Addadi and O. Golani for help in acquisition and analysis of Light Sheet images, and S. Streichan for support in 3D image processing. Imaging using the Light Sheet microscope was made possible thanks to The de Picciotto-Lesser Cell Observatory in memory of Wolf and Ruth Lesser. We thank the members of the Shilo and Barkai labs for fruitful discussions. The work was supported by an ERC advanced grant to N.B. and a US-Israel Binational grant to E.S. and B.S. B.S. is an incumbent of the Hilda and Cecil Lewis chair in Molecular Genetics.

## Methods

### Fly stocks and genetics

For the *wt* dlGFP experiments we used Sco/Cyo;dl-GFP/Tm3, Sb flies (DeLotto et al., 2007). For the *wntD* mutant background, a recombination between a *wntD* null allele (Rahimi et al., 2016)and *dl-GFP* was carried out, and crossed to *wntD* mutant.

*wntD::MS2-Fhos-RH* generation: the 1162 bp upstream of *wntD* transcription start site were synthesized followed by 24 repeats of the MS2 sequence, and placed within the 5UTR. The sequence was inserted into a pAttB vector with *NotI* and *KpnI* sites. The *Fhos-RH* sequence was further ligated into the *NotI* site to generate wntD::MS2-Fhos-RH. Virgin females expressing both MCP::GFP and His:RFP were crossed with males of the reporter line *wntD::MS2-Fhos-RH* to collect embryos for imaging.

MCP::GFP and *t4:MS2-yellow* fly lines (Lim et al., 2017) used in the quantitative live imaging of *T48* induction were kindly provided by B. Lim. Virgin females carrying both MCP::GFP and His:RFP were crossed with males of the reporter line *T48::MS2-yellow* and embryos collected for imaging.

#### Live Imaging

Embryos were imaged using a Light Sheet z1 microscope (Zeiss Ltd.) equipped with 2 sCMOS cameras PCO-Edge, 10X excitation objectives and Light Sheet Z.1 detection optics 20×/1.0 (water immersion). The embryos were collected, dechorionated and up to 4 embryos were sequentially mounted perpendicularly into a glass capillary (Brand) in a 1% low melting agarose solution (Roth). Imaging was preformed using dual side illumination, zoom X0.8. GFP Excitation: 488nm Emission / detection – BP 505-545, RFP Excitation: 561nm Emission / detection – BP 575-615.

The *T48::MS2* expressing embryos were mounted on a cover slide and imaged through halocarbon oil in a Zeiss LSM710 confocal system at a temporal resolution of 3 minutes.

#### MS2 analysis

MCP-GFP spots were manually counted using the Imaris software after adjusting the contrast min&max for enhanced visualization. In the *T48::MS2* analysis the spots were manually detected and displayed using the Imaris spots object.

#### sm-FISH

Stellaris RNA FISH probe sets for the *T48* gene (5’ UTR and coding, not including the 3’ UTR), were designed by Stellaris Probe Designer and purchased from LGC Biosearch Technologies. 3 hrs after egg lay (AEL) *wt* embryos were fixed for 25 min in 4% formaldehyde, washed in Methanol and kept at −20°C. Next day embryos were washed in Methanol and then in Ethanol, rocked in 90% Xylene, 10% Ethanol for 1 hr followed by post fixation. Then incubated 6’ with Proteinase K and post fixed again. Embryos were transferred gradually to 10% FA in 2X SSC + 10 μg/ml ssDNA preheated to 37°C and prehybredized for 30’ at 37°C. Hybridization Buffer included 10% FA, 10% Dextran, 2mg/ml BSA, RVC and ssDNA+ tRNA in 2X SSC, containing the probe set (1 ng/μl) (Trcek et al., 2017). Hybridization was carried out O/N at 37°C. Next morning the embryos were shaken gently and incubated for another 30’. Embryos were washed twice for 30’ at 37°C with 10% FA in 2X SSC + 10 μg/ml ssDNA and gradually transferred to PBS-0.5% Tween and mounted with Vectashield+DAPI Mounting Medium (Vector Laboratories Inc.). Fluorescence was visualized with a Nikon Eclipse Ti2 microscope, and analyzed by the TransQuant script as was previously published (Bahar Halpern and Itzkovitz, 2016). TS Intensities were measured via ImageJ.

### Light Sheet movies analysis

#### Projection to 2D

To enable quantitative analysis of the nuclear Dl gradient, we projected the 3D scans of the embryo from the light-sheet microscope, into a 2D flat surface, for every time point imaged. This was possible, since all the nuclei are arranged on the surface of the embryo, whose shape resembles an ellipsoid. This ellipsoid can be projected into a 2D surface, which contains all the nuclei and therefore the entire nuclear Dl gradient. To this end, we used an area preserving transformation with minimal distortion far from the Anterior and Posterior poles, implemented by the IMSANE tissue cartography tool (Heemskerk and Streichan, 2015). IMSANE was used with the following specifications: Planar Illastik surface detector and cylinder chart type. Surface detection was performed on the last time point for each embryo, and the detected surface was then used to project all earlier time points. Since the embryo is, to a good approximation, a cylinder apart from the anterior and posterior poles, embryo circumference was defined as the largest circumference of the ellipsoid fitted to the embryo surface by IMSANE.

#### Nuclei segmentation

The nuclei were detected separately for each time point, using the following segmentation method:

1. Automated local thresholding of the image in order to create a binary mask. Done in ImageJ using the Bernsen algorithm with a contrast threshold of 15.
2. The resulting binary mask underwent further refinement to segment the nuclei using MATLAB image analysis filters:

a. All connected objects in the mask large enough to be nuclei (over 50 pixels in size) were located and classified into 3 size groups: *small* 50-150 pixels, *medium* 150-600 pixels and *large* 600 pixels and over.
b. Each size group underwent erosion using *imerode* and then dilation using *imdilate* with appropriate filter sizes for each group.
c. Objects belonging to the *large* group underwent another round of erosion and dilation with same filter size as first round.
3. The resulting objects in the binary mask were filtered by size to exclude objects too small (under 50 pixels) or too large (over 3000 pixels) to be a single nucleus.
4. Nuclei locations were detected by overlaying the mask on the original image.

#### Measuring the nuclear Dl gradient around the A-P midline

Gradient measurement was performed by first manually discarding all time points between the nuclear cycles. Then, for every detected nucleus, for all remaining time points, the value of nuclear Dl was calculated as the mean intensity inside the nucleus (located by the above segmentation method). The location of the A-P midline was manually selected for each embryo. A rectangular area around the A-P midline was then defined. The width of the area along the A-P axis was 15% of entire A-P length and it spanned the entire D-V axis. For NC14, this definition corresponds to ~8-10 columns of nuclei closest to the A-P midline. Only the nuclei inside this area were taken into account for gradient measurement, monitoring several columns of nuclei along the A-P axis gave rise to averaging of Dl-nuclear intensity along this small window. The spatial axis for the gradient was defined as a relative axis-*x/L,* indicating location on the D-V axis-*x* divided by embryo circumference-*L.* In order to assign a location on this relative D-V axis for each nucleus, the location of the D-V midline was manually selected and defined as *x/L =* 0. This resulted in the raw intensity function, *Dl_raw_(x/L,t)* measuring nuclei intensity along the relative D-V axis, over time. This function, *Dl_raw_(x/L, t)*, was then smoothened **in space, for each time point separately** using the MATLAB *smooth* function with a smoothing coefficient of 0.23. The smoothened data was then fitted by a smoothing spline using the MATLAB *fit* function and evaluated on a 1000 linearly spaced *x/L* locations. The resulting function, *Dl_smooth_(x/L, t),* was plotted in main text figures.

#### Dorsal gradient as a function of time, at specific locations along the D-V axis

For the calculation of Dl-nuclear intensity over time at a specific location 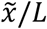, we used *Dl_smooth_(x/L,t)* at that location: 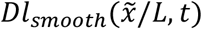. Background subtracted values were calculated separately for each NC, by subtracting the minimal intensity observed in a nucleus for that NC. 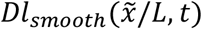 was then smoothened in time using the MATLAB *smooth* function with the *loess* method and a smoothing coefficient of 0.5. It was then fitted with a smoothing spline, resulting in the function 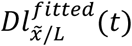. The temporal derivative of nuclear Dorsal, at a specific location-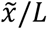 was calculated by applying a third order finite differences formula to 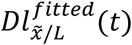 and then smoothing using *smooth* with a smoothing coefficient of 0.6 and fitting a smoothing spline.

#### Measuring peak sharpness

The peak sharpness measure for an embryo was calculated based on the values of *Dl_smooth_(x /L,t)*, around the ventral-most location. For each time point, 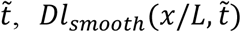 was normalized by dividing by its maximal value at 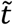. Peak sharpness was then calculated as the standard deviation divided by the mean, in percent, of values close to the peak: within the range 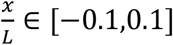. This measure captures how different from each other are values close to the peak.

## Supplemental Information

### Supplemental Movies, figures and Tables

**Movie 1** – A Light Sheet time-lapse movie following the dynamics of endogenously expressed Dl-GFP in the entire embryo. The nuclear Dl gradient can be seen in nuclei at the ventral side (bottom) already at NC 12, it is lost during nuclear divisions and is re-generated at the onset of each nuclear cycle.

**Movie 2** – A frame by frame 2D projection of movie 1 done using the ImSAnE tool (Heemskerk and Streichan, 2015). Dl-GFP appears in grey.

**Movie 3** – Time lapse of Dl-GFP intensity data for the area inside the dashed frame in Figure 1C. Each circular marker in the movie shows raw, non smoothed Dl-GFP intensity in a single nucleus. Nuclei were segmented from the corresponding frames of movie 2 (See Methods).

**Figure S1.**
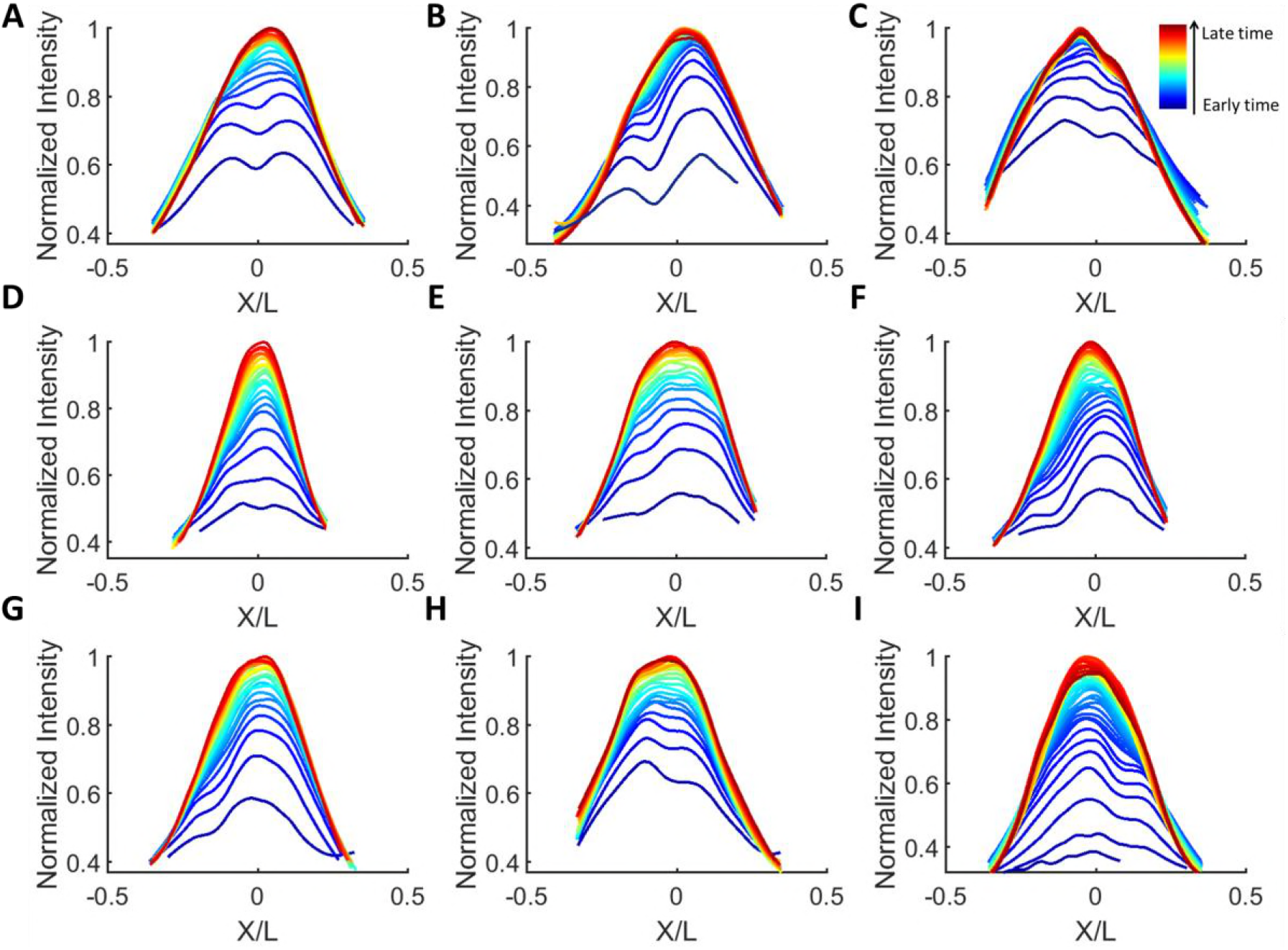
Dl temporal dynamics during NC14 in *wt* embryos. Dl-GFP intensity, plotted as function of relative location along the DV axis, relative location axis x/L is defined as location divided by embryo circumference (See Methods). A curve is shown for each time point during NC14 for 9 *wt* embryos. Each Dl-GFP intensity curve was smoothened and normalized by the maximal value attained during NC14 (See Methods). Time points are 1.5 minutes apart for (A-H) and 1 minute apart for I, going from earliest time points in blue to the latest in red. All embryos exhibit shuttling signatures: lateral overshoots and converging double peaks.

**Figure S2.**
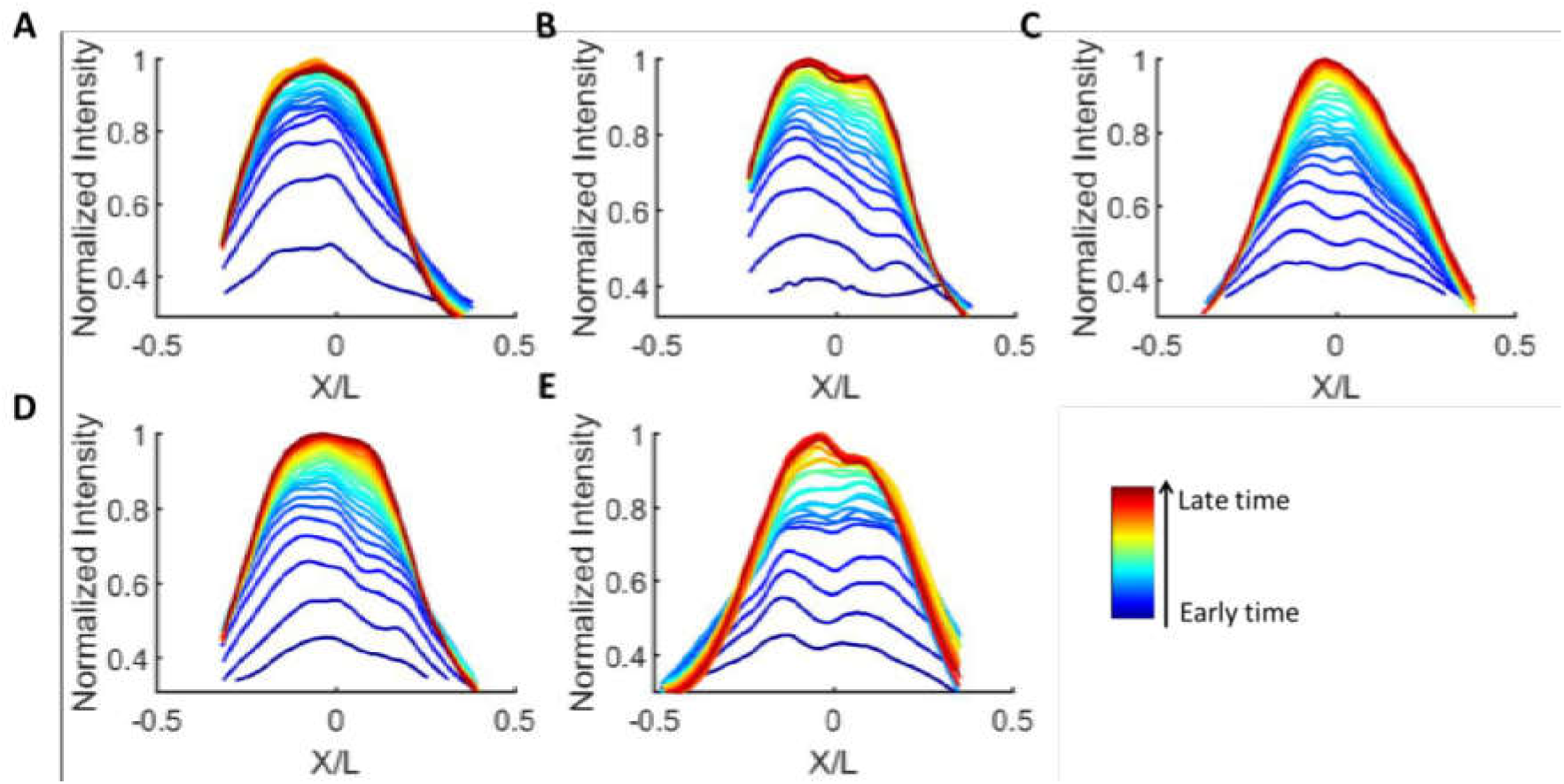
Dl temporal dynamics during NC14 in *wntD-/-* embryos. Dl-GFP intensity, plotted as function of relative location along the DV axis, relative location axis x/L is defined as location divided by embryo circumference (See Methods). A curve is shown for each time point during NC14 for 5 *wntD-/-* embryos. Each Dl-GFP intensity curve was smoothened and normalized by the maximal value attained during NC14 (See Methods). Time points are 1.5 minutes apart for, going from earliest time points in blue to the latest in red. All embryos exhibit shuttling signatures: lateral overshoots and double peaks flattening over time.

**Figure S3.**
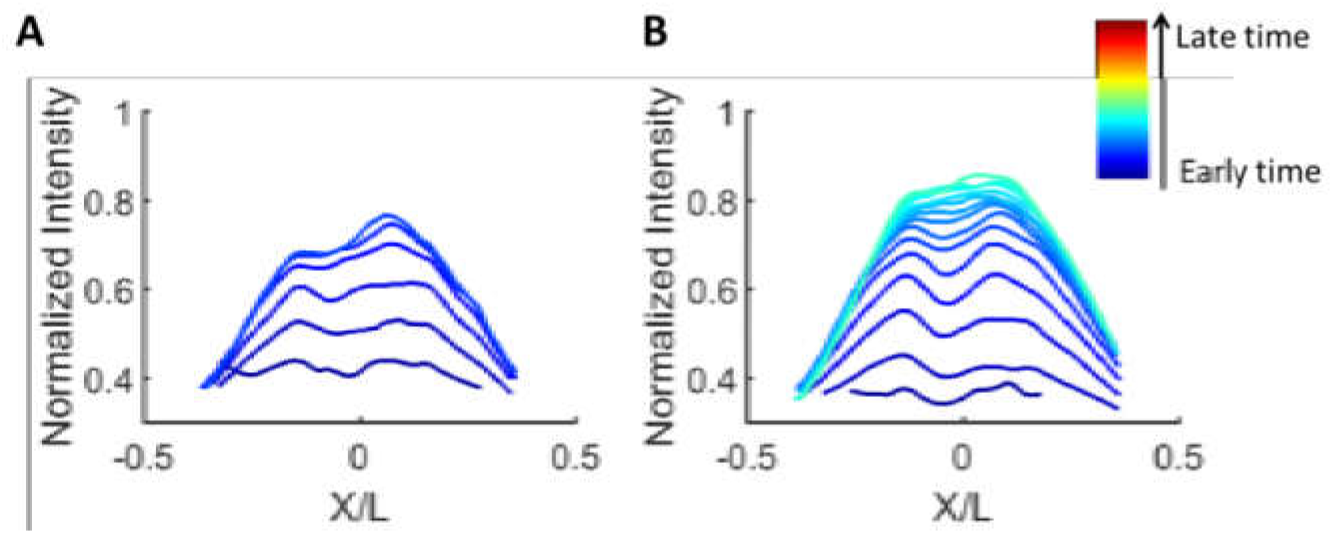
Dl temporal dynamics during NC12-13 in a *wntD-/-* embryo. Dl-GFP intensity, plotted as function of relative location along the DV axis, relative location axis x/L is defined as location divided by embryo circumference (See Methods). A curve is shown for each time point during NC12 (A) and NC13 (B) for the *wntD-/-* embryo presented in Figure 3E. Each Dl-GFP intensity curve was smoothened and normalized by the maximal value attained during NC14 (See Methods, Figure 3E). Time points are 1 minute apart for, going from earliest time points in blue to the latest in red.

**Table S1:** Parameters table for full model, WT values

| # | Parameter | Units type | Meaning | Value in a.u | Units | Value in units |
| --- | --- | --- | --- | --- | --- | --- |
| 1 | $K_{rec}$ | $t^{-1}$ | Toll receptor recycling rate | 0.07 | $sec^{-1}$ | 0.007 |
| 2 | $K_{end}$ | $t^{-1}$ | Toll receptor endocytosis rate | 1 | $sec^{-1}$ | 0.1 |
| 3 | $\lambda$ | $t^{-1}$ | NCSpz* cleavage rate | 4.2 | $sec^{-1}$ | 0.42 |
| 4 | $\alpha_N$ | $t^{-1}$ | NSpz degradation rate | 0.0011 | $sec^{-1}$ | 0.00011 |
| 5 | $\alpha_c$ | $t^{-1}$ | CSpz degradation rate | 0.0014 | $sec^{-1}$ | 0.00014 |
| 6 | $\alpha_{Nc}$ | $t^{-1}$ | NCSpz degradation rate | 0.0016 | $sec^{-1}$ | 0.00016 |
| 7 | $K_{on,c}$ | $t^{-1}C$ | CSpz binding to Toll | 0.3 | $\mu Msec^{-1}$ | 0.0013 |
| 8 | $K_{off,c}$ | $t^{-1}C$ | CSpz un-binding from Toll | 1.12 | $\mu Msec^{-1}$ | 0.0049 |
| 9 | $K_{on}$ | $t^{-1}C$ | NCSpz binding to Toll | 0.008 | $\mu Msec^{-1}$ | 3.2e-5 |
| 10 | $K_{off}$ | $t^{-1}C$ | NCSpz un-binding from Toll | 0.1 | $\mu Msec^{-1}$ | 4.33e-04 |
| 11 | $\eta_0$ | $t^{-1}C$ | NCSpz activation rate | 0.22 | $\mu Msec^{-1}$ | 9.68e-4 |
| 12 | $K_{bind}$ | $t^{-1}C$ | CSpz binding rate to NSpz | 7 | $\mu Msec^{-1}$ | 0.0308 |
| 13 | $K_{split}$ | $t^{-1}$ | NC-Tl splitting rate to C-Tl and 2*N-Spz , on Toll | 10 | $sec^{-1}$ | 1 |
| 14 | $D_{NC}$ | $x^2t^{-1}$ | NCSpz diffusion coefficient | 0.1186 | $\mu m^2sec^{-1}$ | 118.6 |
| 15 | $D_N$ | $x^2t^{-1}$ | NSpz diffusion coefficient | 0.0055 | $\mu m^2sec^{-1}$ | 55 |
| 16 | $D_{NC*}$ | $x^2t^{-1}$ | NCSpz* diffusion coefficient | 0.745 | $\mu m^2sec^{-1}$ | 745 |
| 17 | $T_{tot}$ | $C$ | Total Toll level | 5.625 | $\mu M$ | 0.25 |
| 18 | $D_W$ | $x^2t^{-1}$ | WntD diffusion coefficient | 0.5 | $\mu m^2sec^{-1}$ | 500 |
| 19 | $\alpha_w$ | $t^{-1}$ | WntD degradation rate | 0.001 | $sec^{-1}$ | 1.4e-5 |
| 20 | $\beta_w$ | $t^{-1}C$ | WntD production rate | 0.1 | $\mu Msec^{-1}$ | 4.4e-4 |
| 21 | $K_w$ | $C^{n_w}$ | $K_D$ of WntD production as a function of signaling | 9e-10 | $\mu M^{n_w}$ | 4.7e-30 |
| 22 | $n_w$ | - | Hill coefficient of WntD production as a function of signaling | 15 | - | 15 |
| 23 | $L$ | $x$ | Length of tissue | 2 | $\mu m$ | 250 |
| 24 | $X_{pipe}$ | % | % of L below which a point $\vec{x} = (x, y, z) \in pipe\ domain$ | 40 | % | 40 |
| 25 | $X_{WntD}$ | % | % of L above which a point $\vec{x} = (x, y, z) \in WntD\ producing\ domain$ | 25 | % | 25 |
| 26 | $W_{lower}$ | % | Maximal lowering of Tl property by WntD | 60 | % | 60 |
| 27 | $W_{Tr}$ | $C^{n_{lower}}$ | $K_D$ of WntD lowering receptor property | 5 | $\mu M^{n_{lower}}$ | 0.01 |
| 28 | $n_{lower}$ | - | Hill coefficient of WntD lowering receptor property | 2 | - | 2 |
| 29 | $T$ | $t$ | # of discrete time points the simulation was run | 600 | $sec$ | 6000<br>(~1.5 hours) |
| 30 | $K_{Dl,in}$ | $t^{-1}C$ | Rate of Dl entry into the nucleus | 0.07 | $\mu Msec^{-1}$ | 2.8e-04 |
| 31 | $K_{Dl,out}$ | $t^{-1}C$ | Rate of Dl exit from the nucleus | 2 | $\mu Msec^{-1}$ | 0.008 |
| 32 | $Dl_0$ | $C$ | Initial Dl concentration in the nuclei | 1e-4 | $\mu M$ | 4.4e-06 |
| 33 | $Dl_{tot}$ | $C$ | Total amount of Dl | 5 | $\mu M$ | 0.22 |

### Mathematical model

Our full model describes how the Spz gradient is formed by shuttling, induces the nuclear localization of Dl and the negative feedback between WntD and Dorsal which maintains the gradients robustness (Rahimi et al., 2016). To this end, we extend the model from our previous paper (Rahimi et al., 2016) to include the nuclear localization of Dorsal. Also, we used a different mechanism by which WntD contracts the Spz gradient: instead of competing with the ligand for binding the Toll receptor, we assume here that WntD “crowds” the Toll receptors immediate environment by binding its own receptor Frizzled4 and limiting the access of Spz to Toll. This increases the chances of free ligand binding the shuttling molecule instead of the receptor and therefore enhances shuttling. This “crowding” of Toll has an additional affect: stabilizing the ligand which succeeded in binding, which also makes shuttling more efficient. The governing set of reaction-diffusion equations of our model is given below: **eqn. 1** defines the temporal dynamics of freely diffusing WntD. The terms of the equation by order of appearance describe: WntD diffusion, WntD degradation, WntD production which depends on nuclear Dorsal. This last term is the induction part of InC as WntD production is positively regulated by signaling. The WntD producing zone is restricted and is defined in the embryo by the Torso signaling border. In the simulations we define this zone using the model parameter *X_WntD_*: WntD production is only allowed for points posterior to *X_WntD_*. **Eqn. 2** defines the nonlinear saturating function 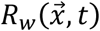 through which WntD changes the properties of the Toll receptor: the rate at which ligands bind and unbind from it. **Eqn. 3** defines the temporal dynamics of free Toll receptors. This equation introduces the following constraint: the total amount of Toll receptors (free, bound by ligand and endocytosed) is constant and equals *T*_tot_. **Equations 4–10** are the self-organized shuttling model (SOSH) equations as appear in (Haskel-Ittah et al., 2012), we’ll review SOSH and the equations briefly. The SOSH mechanism depends on the versatility of the *Spz* protein. The separate inhibitor domain N-Spz and activating region of *Spz,* C-Spz, generated after cleavage of the NC-Spz precursor (**eqn. 4**), can interact with each other in three different modes. These modes facilitate a process of “selforganized shuttling”, where the active ligand C-Spz is shuttled and concentrated at the ventral-most region giving rise to the sharp activation gradient of *Toll.* **Equations 5–7** describe the shuttling of the active C-Spz ligand (which cannot diffuse on its own) by the N-Spz inhibitor when bound together as the NC-Spz* complex. Signaling occurs when C-Spz bound to Toll undergoes endocytosis (**eqns. 5,8,10**). Toll receptors undergo recycling back to the membrane after endocytosis and the total concentration of Toll is constant (**eqns. 10,3** respectively). The C-Spz ligand and N-Spz inhibitor are products of NC-Spz complex separation when bound to the Toll receptor (**eqn. 4,9**). NC-Spz is also capable of inducing Toll endocytosis when binding it and thus contributing to signaling (**eqn. 9**) but signaling through NC-Spz happens at a much lower rate than C-Spz mediated signaling. **Equations 11–12** describe the induction of Dorsal nuclear localization by Toll signaling. **Eqn. 11** introduces the following constraint: the total amount of Dorsal (Nuclear-*Dl_in_* and cytoplasmic-*Dl*_out_) is constant and equals *Dl*_tot_. The meaning of the different parameters and their units are summarized in Table S1. This set of equations was solved numerically in 1D using a standard MATLAB PDE solver.

#### Full model equations

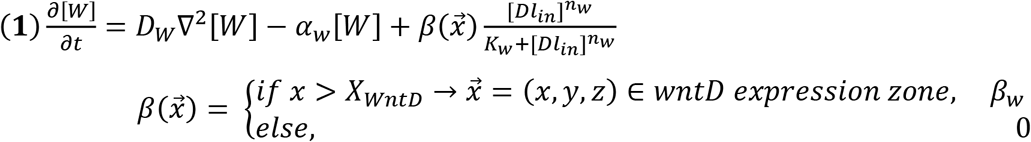

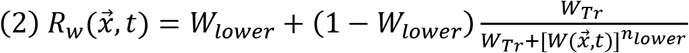

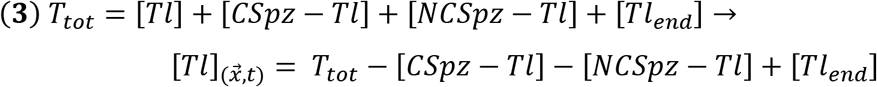

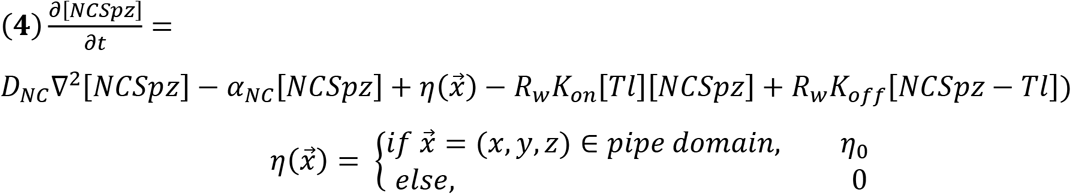

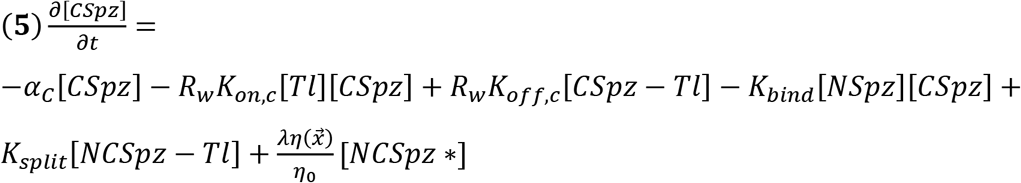

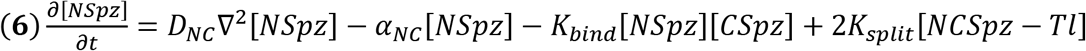

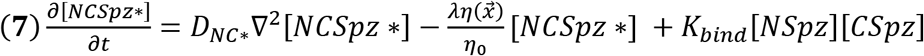

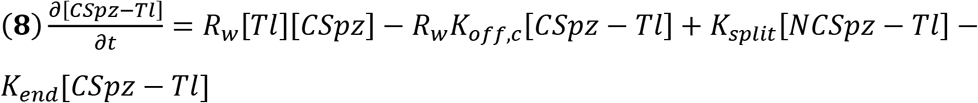

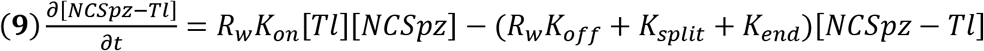

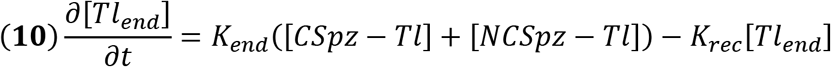

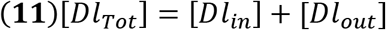

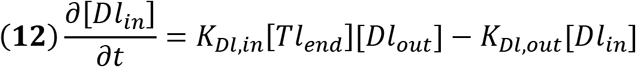

#### Full model parameters

When choosing parameters, we relied on our previous papers where SOSH (Haskel-Ittah et al., 2012) and InC (Rahimi et al., 2016) were previously analyzed. For the SOSH WT parameters we in this work, we used the SOSH parameter values described in Table S5 of (Haskel-Ittah et al., 2012) with slight alterations to better resemble our experimental data. Values for WntD related parameters (diffusion, degradation, and production rates) were based on our previous paper (Rahimi et al., 2016). The addition of Dorsal nuclear localization (eqns. 11–12) introduced new dorsal related parameters, which were selected to resemble similar parameters in the model (for example, the total concentration of Dorsal is similar to that of Toll). For the simulation of WntD mutants, model equations were solved with the WntD production rate set to zero.

##### Constant external gradient model

In order to simulate only Dorsal nuclear localization induced by a constant Toll signaling profile, the following set of equations was solved:

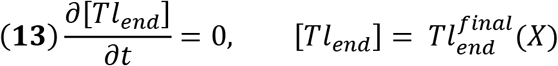

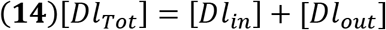

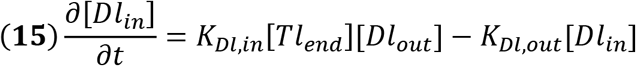

The constant external gradient, 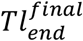, was selected to be the sharp Toll signaling profile [*Tl_end_*] at the last time point for the simulation of the full model (eqns 1–12). Parameter values according to Table S1.

#### Naïve diffusion model

This model includes a single ligand, *NCSpz,* which is produced throughout the pipe domain (eqn. 17). The ligand diffuses and is degraded (eqn. 17). It binds the Toll receptor (eqns. 18,16), and induces Dorsal nuclear localization (eqns. 20–21). This model is described by the following equations:

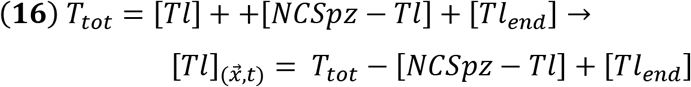

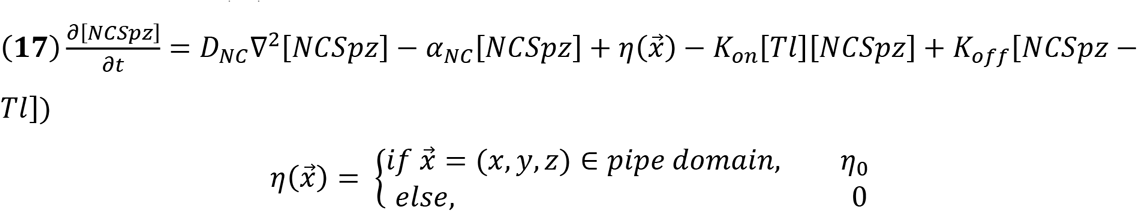

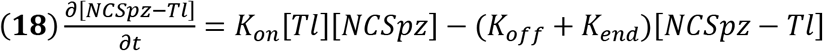

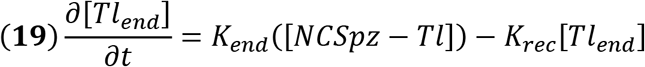

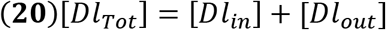

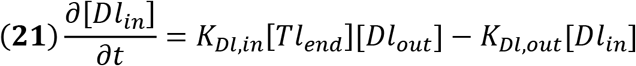

Parameter values according to Table S1. See further analytical and numerical analysis for the sharpness and robustness of the gradient in this model in (Haskel-Ittah et al., 2012).

#### Shuttling parameters effect on double peak prominence

The prominence of the converging double peak feature in the shuttling model depends on model parameters which control the mean path the shuttling complex travels ventrally before being cleaved, relative to the length of the source (pipe domain). This mean path is influenced by several parameters, mainly tissue absolute length *L* (when maintaining the source as 40% of absolute length), changing source length directly by assuming the pipe domain is smaller or larger than 40% of the circumference, cleavage rate of the shuttling complex *λ*, diffusion coefficient of the shuttling complex *D_NC*_*. For a sufficiently large ratio between the mean path and *L*, no double peak will be observed. Lowering this ratio increases the prominence of the double peak. For a sufficiently small ratio, the double peak does not converge into a single peak since the shuttling complex cannot penetrate all the way to the ventral most. We demonstrate this by solving the full model for our WT parameter set (Table S1), and comparing to sets where these parameters are perturbed: decreased 2 fold and increased 2 fold:

**Model Supplement Figure 1.**
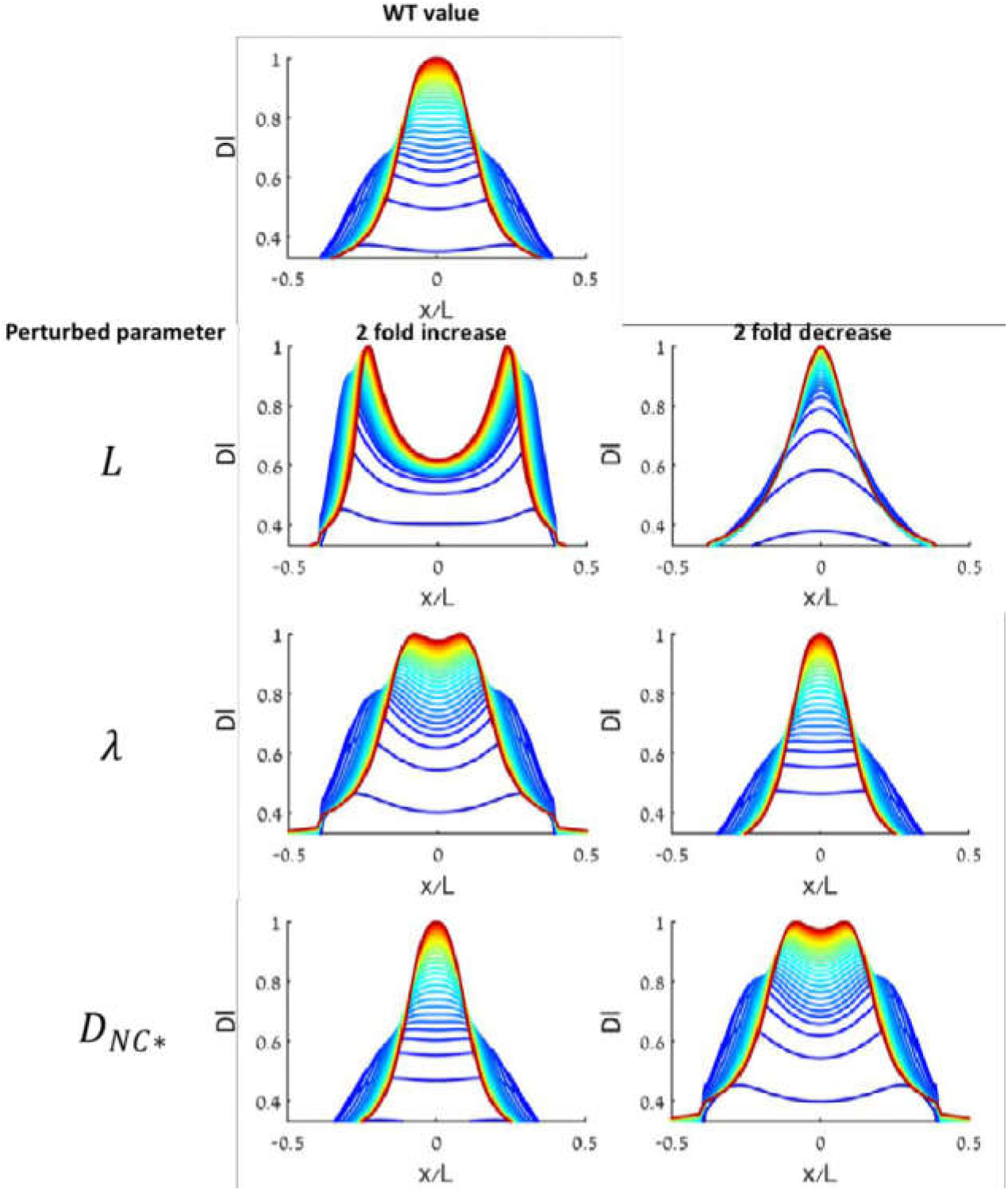
Perturbations to shuttling complex parameters modulate double peak prominence.

